# Migration of human mesenchymal stem cells stimulated with pulsed electric field and the dynamics of the cell surface glycosylation

**DOI:** 10.1101/122382

**Authors:** Katarzyna Jezierska-Wozniak, Seweryn Lipiński, Łukasz Grabarczyk, Monika Barczewska, Aleksandra Habich, Joanna Wojtkiewicz, Wojciech Maksymowicz

## Abstract

The objective of our study was to develop novel techniques for investigations of cell motility, and to assess whether the electric field of the therapeutic spinal cord stimulation system used *in vivo* contributes to the migration of human mesenchymal stem cells (hMSCs) *in vitro.*

We have investigated electrotaxis of bone marrow-derived MSCs using pulsed electric field (PEF) in range 16-80 mV/mm and frequency 130 Hz and 240 Hz. The PEF-related dynamics of the cell surface glycosylation was evaluated using six plant lectins.

PEF at physiological levels (10mV/mm; 130 Hz) did not influence cellular motility *in vitro*, what may correspond to the maintenance of the transplanted cells at the lesion site *in vivo*. Increase of the PEF intensity and frequency above physiological levels resulted in the increase in the cellular migration rate *in vitro*. PEF elevated above physiological intensity and frequency (40-80 mV/mm; 240 Hz), but not at physiological levels, resulted in changes of the cell surface glycosylation.

We find the described approach as convenient for investigations and for the *in vitro* modeling of the cellular systems intended for the regenerative cell transplantations *in vivo*. Probing cell surface glycomes may provide valuable biomarkers to assess competence of transplanted cells.

## Introduction

Stem cell therapy encompasses new technologies and therapies aiming at replacing damaged cells with healthy ones. However, there are still several challenges that must be tackled before such therapies become successful. One of these challenges is the migration of transplanted cells, fundamental both to the development and maintenance of regenerated tissues as well as to the success of the cell-based therapies. Therefore, precise localization of these cells directly into the injury site and their adaptation to host tissues is crucial for the success of therapy.

Delivering transplanted cells precisely at the lesion site is still an unsolved problem of cell therapy. This process, termed homing, is a major road-block in the stem cell therapy due to the poor integration of transplanted stem cells with a targeted host tissue. This results in poor healing responses and lack of regeneration. The main reason for poor cell homing is likely due to the fact, that only a small fraction of injected cells migrating to the damaged tissue and are able to act therapeutically (Okano and Sawamoto, 2008; Smart and Riley 208; Dai et al., 2005). It is also critical to demonstrate that MSCs do not have unwanted homing that could drive undesirable differentiation and growth (Breitbach et al., 2007).

The cells migrate in response to gradients in chemical composition (chemotaxis), mechanical forces, and electric fields (galvanotaxis or electrotaxis). Electric fields (EFs) are present in all developing and regenerating tissues. Furthermore, damaged tissue and wounds also generate naturally-occurring endogenous EFs essential in guiding cell migration (McCaig et al., 2005). Endogenous EFs have been identified and measured in various species, tissues and organs *in vitro* and *in vivo* (Barker et al.,1982; Levin et l., 2006; Zhao et al., 2006). It has been shown, that many cell types respond to applied EFs *in vitro* at the field strengths comparable to the endogenous wound EFs *in vivo*. The bioelectricity plays an important role in moving cells in to a specific direction. Rat osteoblasts, bovine chondrocytes, and mouse endothelial progenitor cells migrate towards the cathode (Chao et al., 2000; Ferrier et al.,1986; Ozkucur et al., 2009; Zhao et al., 2011), while rabbit osteoclasts, human osteosarcoma cells, rabbit corneal endothelial cells or human umbilical vein endothelial cells migrate in the opposite direction (Ferrier et al.,1986; Ozkucur et al., 2009; Zhao et al., 2004). These different responses indicate that the effects of electric current are cell type- and species-dependent. Consequently, applied electric field may be used as a cue to guide directional migration of stem cells in the cell-based therapies (McCaig and Zhao 1997).

The most common assays for the *in vitro* electrotaxis studies employ cell culture dish attached to the electrotaxis apparatus (Zhao et al., 2006; Sato et al., 2007). Microfluidic devices allowing better control of the electric field and cell migration environments have been recently developed (Li and Lin 2011;Li et al., 2012). New solutions, providing advanced experimental tools for electrotaxis research, improved productivity and enabling studies of the more complex cell migration environments have also been recently reported (Lin et al., 2008; Huang et al., 2009; Huang et al., 2009; Tsai et al., 2012; Zhang et al., 2011; Sun et al., 2012; Schumann et al., 2010 Rezai et al., 2012). However, these experimental tools allow investigations of cellular motility only in the laboratory conditions, making transfer of the favorable experimental findings to the clinical setting difficult.

One of the major macromolecular systems defining cell-cell and cell extracellular matrix interactions is the cell glycome consisting of a large variety of complex glycans covalently attached to the membrane proteins and lipids. These glycoproteins and glycolipids are actively involved in, and often control cell-cell signaling, immune response, would healing, microbial infections, and other events on the cellular and tissue levels. The dynamics of glycomic profile reflects changes in the cellular activities such as division, differentiation, motility, secretory functions and malignant transformation. Cell glycosylation-targeting investigations would therefore be expected to streamline discovery of novel biological factors of potential clinical significance (Alley et al., 2013).

The aim of this study was to develop novel technique for cell motility investigations, and to assess whether the system used for spinal cord stimulation is able to direct the migration of MSCs *in vitro*. We first used pulsed electric field to observe the phenomena of MSCs electrotaxis. Using time–lapse video microscopy, we demonstrated and analyzed directs migration of the MSCs in the pulsed electric field. Additionally, using six plant lectins, we have also evaluated changes in the cell surface glycosylation resulting from the stimulation of cells with pulsed electric field.

## Methods

### Cell culture

We used adult human MSCs derived from the bone marrow remaining after the hip replacement. The experimental design has been approved by the Ethic Committee of University of Warmia and Mazury (UWM) in Olsztyn. Written informed consent was obtained from each participant prior to the study enrollment. MSCs were isolated from the bone marrow according to their adhesive properties to tissue culture plastic under sterile conditions. Briefly, a phosphate-buffered saline (PBS)-diluted cell fraction of bone marrow was layered over a Ficoll density gradient (1.077 g/mL, GE Healthcare), followed by centrifugation at 400G at room temperature for 40 min. Nucleated cells were collected, diluted with two volumes of PBS (Sigma-Aldrich, Saint Louis, USA), centrifuged twice at 100G for 10 min, and finally resuspended in culture medium. Cells were plated at a density of 1,500 cells/cm^2^ in a T75 flask and serially passaged at sub-confluence every 5-7 days at the same initial density. Cells were maintained in the DMEM growth medium supplemented with 10 % (v/v) FBS (Sigma-Aldrich, Saint Louis, USA) and 1 % (v/v) antibiotics 1% 10,000-U penicillin/streptomycin (P/S) (Sigma-Aldrich, Saint Louis, USA), at 37°C in an air-5 % CO_2_ incubator. For the evaluation of the electrical field-dependent changes in the cell surface glycan profiles cells were seeded on the thin glass slides and treated with the electrical field stimulation using parameters described below.

### EFs stimulation and Time Lapse Image Recording

Cells (1,500 cells/ cm2) were seeded and grown for at least 12 h in culture conditions as described above. To test the effect of electric field and to ensure a constant flow of current, in case of the impendence drop, temperature change and other similar technical issues, a pulsed electric field (PEF) was applied through bipolar electrode (St. Jude Medical Inc.). Electric field intensity of 4 mA (16mV/mm), 10 mA (40mV/mm) and 20 mA (80 mV/mm) were used in the experiments, with an exposure times of 3, 6 and 9 hours. Stimulation frequencies were set at 130 Hz and 240 Hz, respectively to the field intensities. The pulse wave was set to 200 microseconds.

Time-lapse imaging was performed using an inverted microscope (JuLi FL; NanoEntek) to digitally record MSC migration. Images were acquired every 15 minutes. For long-term observations, the cell cultures as well as the PEF stimulation device were kept in the air-CO2 incubator. At the end of exposure period, cells were fixed in the PBS-buffered 4% formalin and then digitally photographed for quantification.

### Evaluation of cells movement

Each migrating cell was traced using the position of the centroid of the plane figure, covering the area occupied by the cell in image obtained for each timeframe. Human operator generated outline of each cell manually. The centroid of the cell represents cell’s centre of gravity, with coordinates ( *X* and *Y*) given as (Malina and Smiatacz et al., 2005):

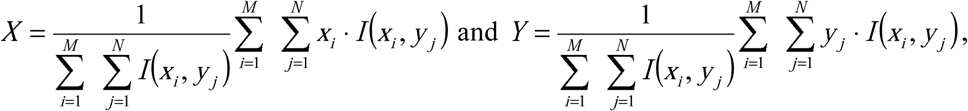

where:

*M*, *N* - image dimensions,

*I*(*x*_*i*_, *y*_*i*_) - value of the pixel with *x*_*i*_, *y*_*i*_ coordinates (in our case each image containeds binary information and so *I*(*x*_*i*_, *y*_*i*_) equals 1 when pixel of these coordinates represents area covered by the cell, otherwise it is equal to 0).

Using the centre of gravity to localize so called "blob" (i.e. figure of an irregular shape) in a binary image is a commonly used approach (Batchelor and Waltz 2012) and can be successfully utilized in experimental applications (Zielinski and Strzelecki 2002). Figure 1 shows the centroid obtained for sample cell. Based on the locations of the cells in each timeframe, the parameter of distance was calculated section-by-section and as the straight-line distance between the first and the last frame, in order to obtain information about the linearity of migration, which can be evaluated through the straightness factor, i.e. the ratio of the straight line distance to the total distance. All measurements were made in triplicates.

**Fig.1.**
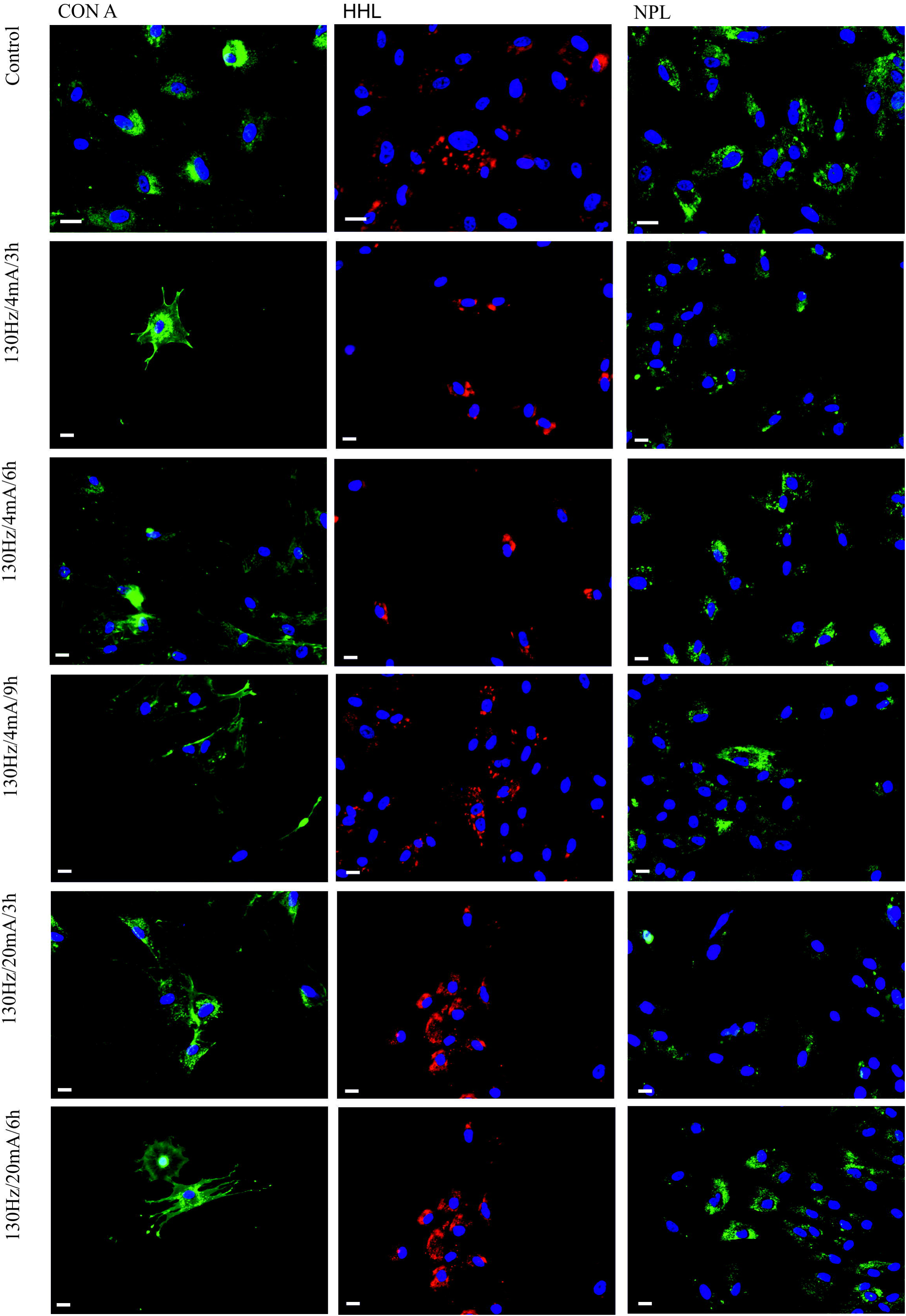
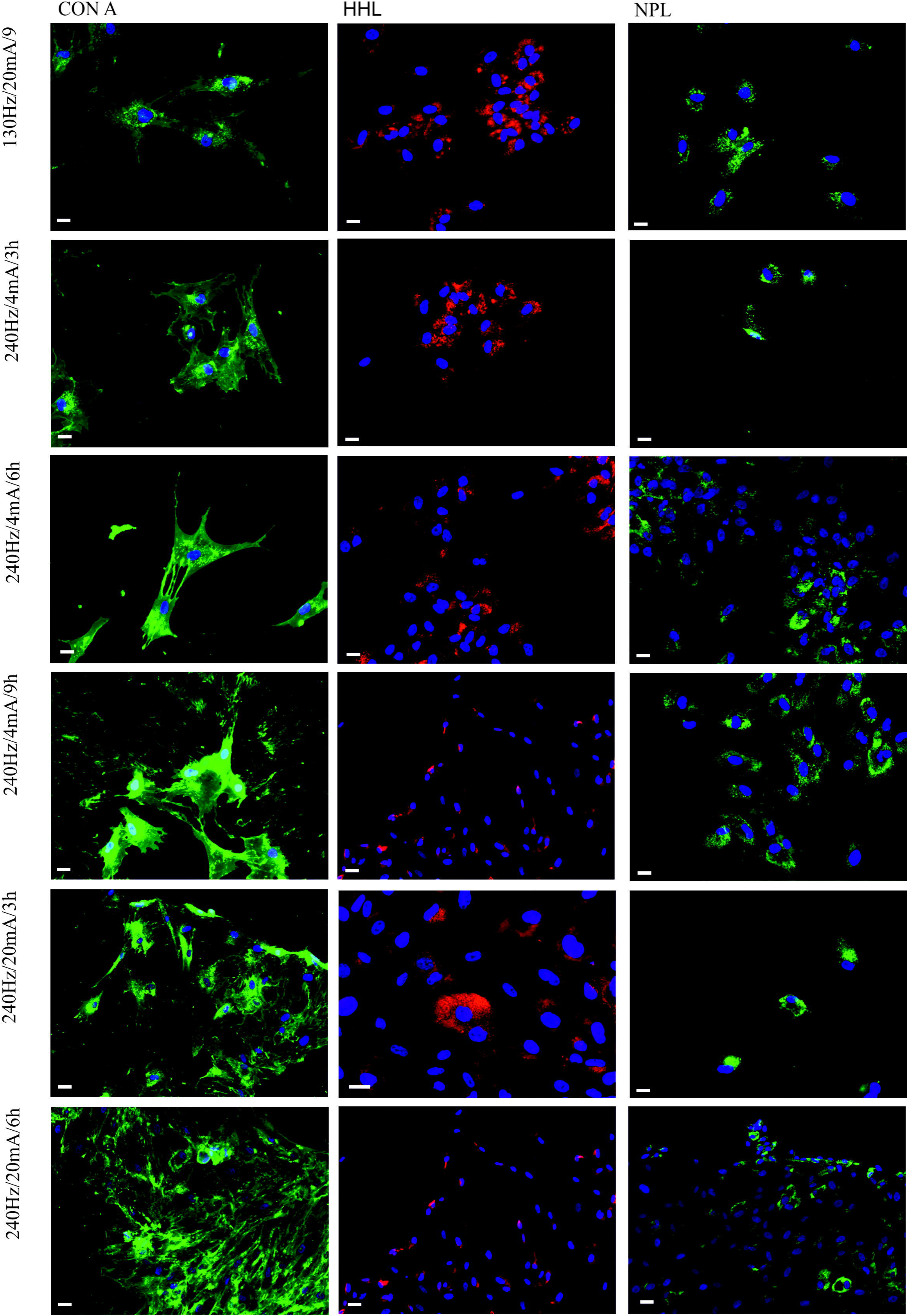
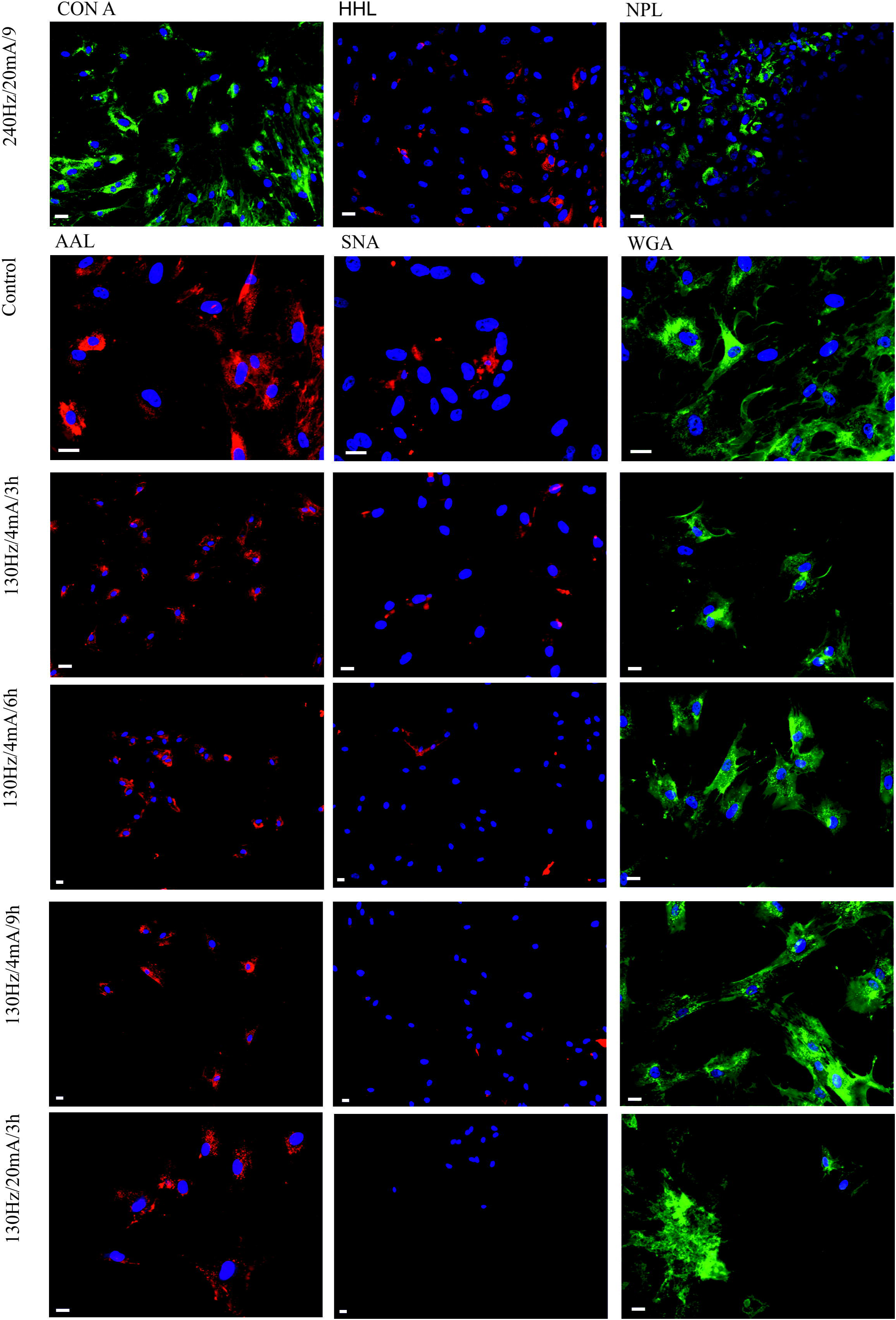
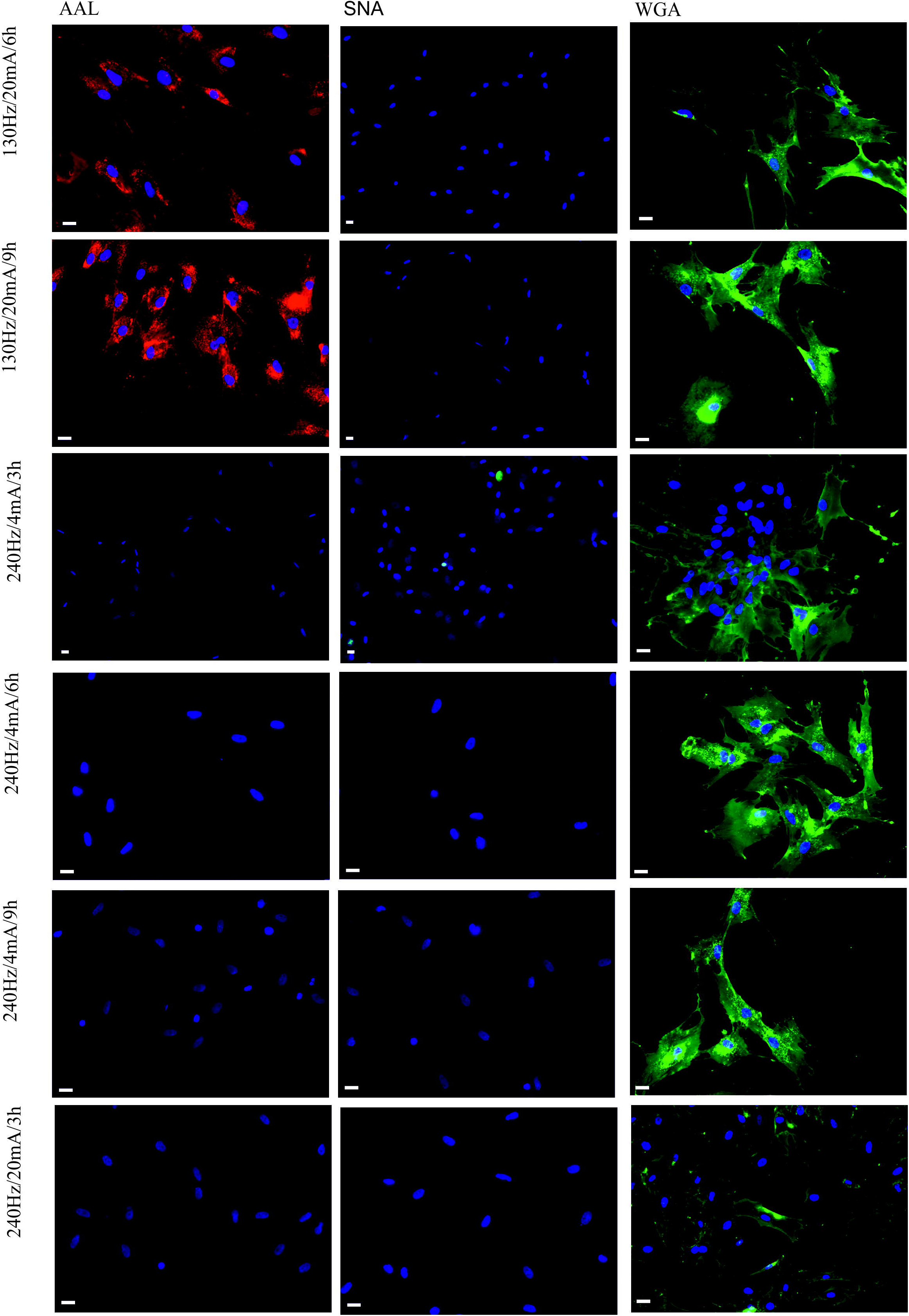
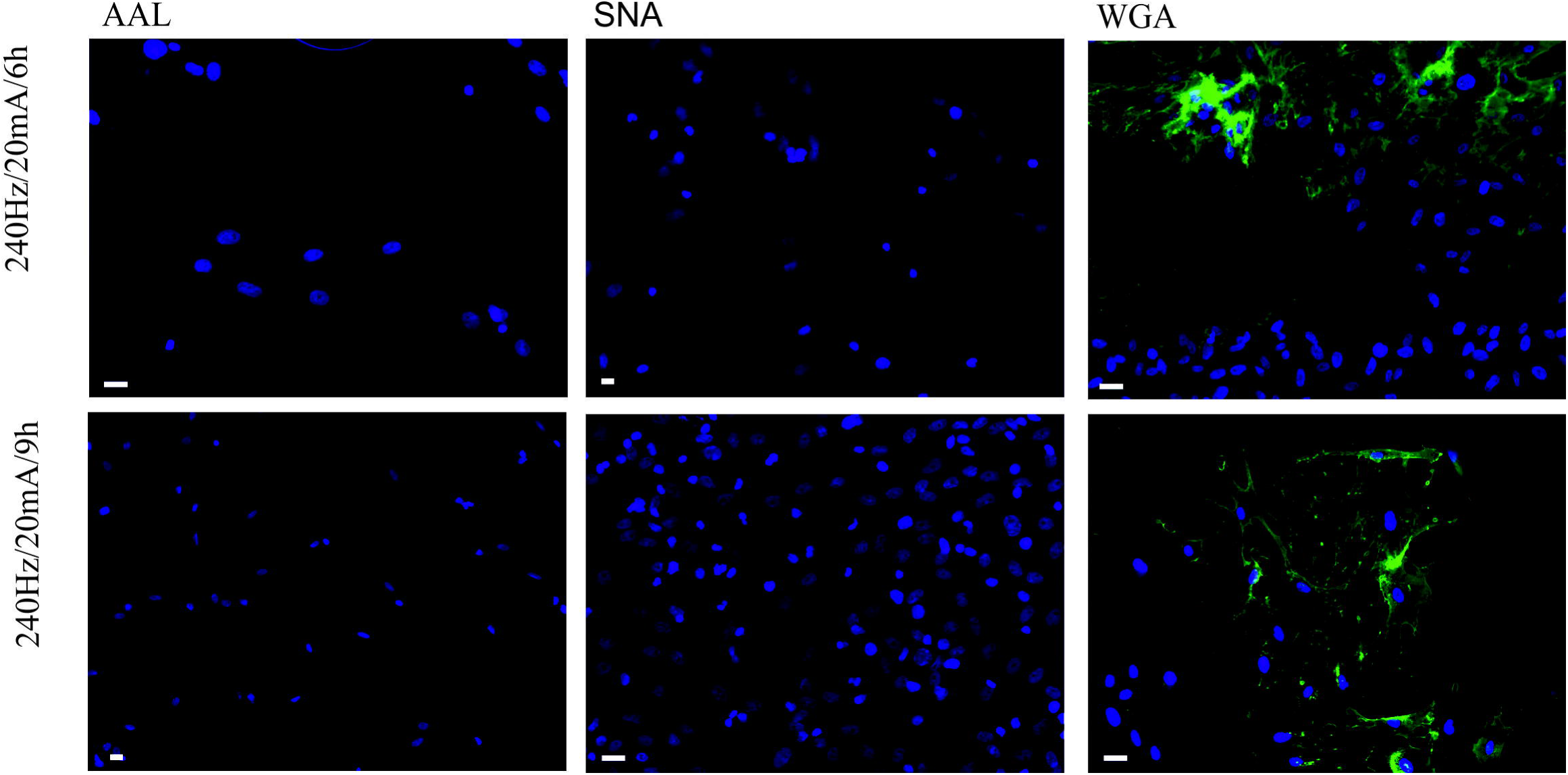
Centroid of the exemplary cell (indicated by the intersection of the grey lines).

### Cell viability

Cell survival was assessed using a commercial assay The CellTiter 96 AQueous One Solution Cell Proliferation Assay (Promega), according to the manufacturer's protocol.

### Lectin binding

Following PEF treatment, cells were washed twice with PBS and fixed immediately for 15 minutes in the PBS-buffered 4% formalin. The carbohydrate phenotype was analyzed using six biotinylated lectins of plant origin: *Aleuria Aurantia* (AAL), *Hippeastrum Hybrid* (Amaryllis) (HHL), *Sambucus Nigra* (SNA), *Narcissus Pseudonarcissus* (NPL), Wheat Germ Agglutinin (WGA), and Concanavalin A (CON A) (Table I). All lectins were purchased from Vector Labs (San Diego, USA). Fixed cells were incubated with 2 μg/mL of each biotinylated lectin overnight, washed three times with PBS and incubated with FITC- Streptavidin (Invitrogen) and Cy3- Streptavidin (Invitrogen), in 1:2000 dilution. Cellular nuclei were stained with the Hoechst 33258 dye (Sigma-Aldrich, Saint Louis, USA).

**Table 1.**
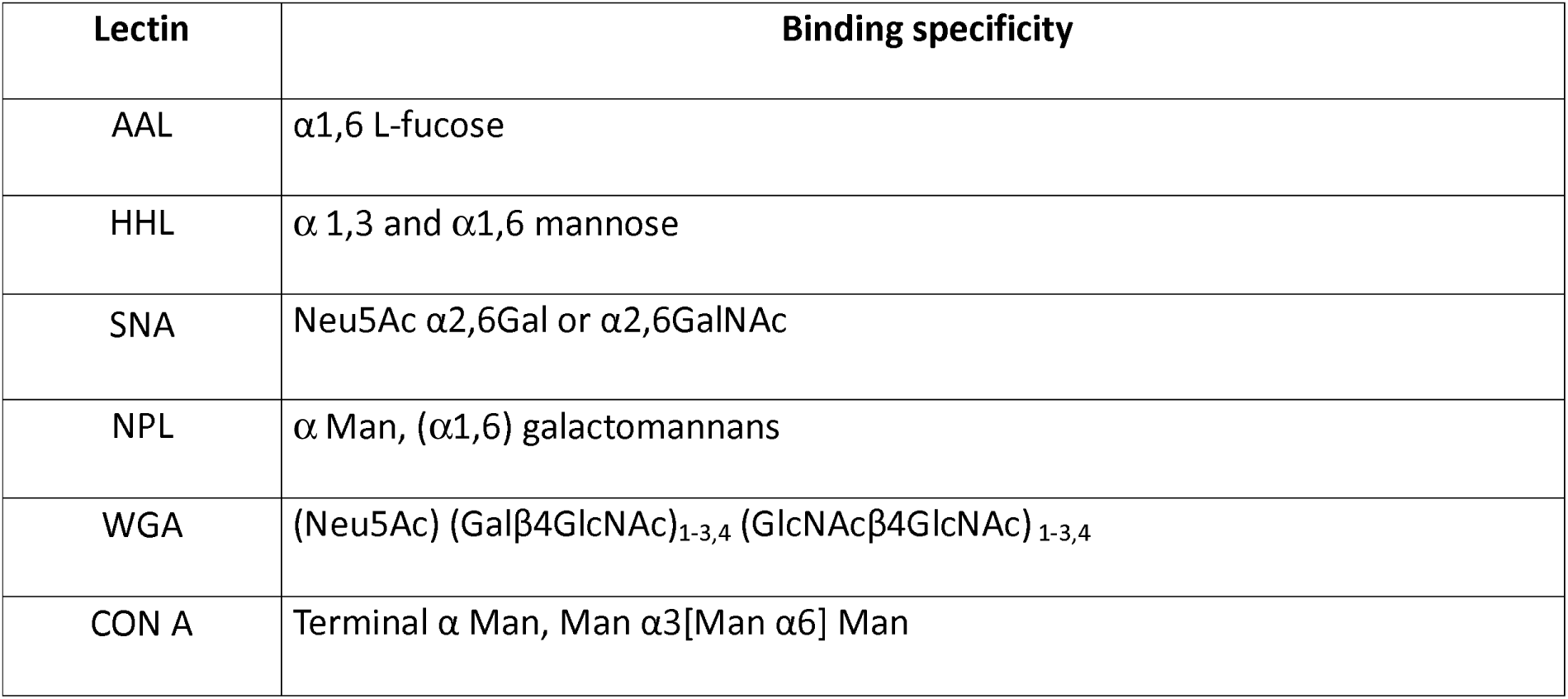
Lectins used in this study and their glycan-binding specificity.

| Lectin | Binding specificity |
| --- | --- |
| AAL | α1,6 L-fucose |
| HHL | $\alpha$ 1,3 and α1,6 mannose |
| SNA | Neu5Ac α2,6Gal or α2,6GalNAc |
| NPL | $\alpha$ Man, (α1,6) galactomannans |
| WGA | (Neu5Ac) (Galβ4GlcNAc)<sub>1-3,4</sub> (GlcNAcβ4GlcNAc)<sub>1-3,4</sub> |
| CON A | Terminal $\alpha$ Man, Man α3[Man α6] Man |

## Results

### Evaluation of cellular movement

The experiment was conducted in two steps. The first step aimed to determine whether the frequency of the pulsed electric field had an impact on the cell movement. As our preliminary results indicate, cellular motility increases with the value of the electric field (Jezierska-Wozniak et al., 2016), so we used two frequencies 130 and 240 Hz, of the highest value of field intensity, i.e. 80 mV/mm, with a 3 hours of observation time. We evaluated statistical differences in the total distance (i.e. distance measured section by section), as well as in the straight-line distance. The Kruskal-Wallis test with the 0.01 significance level showed that there is no significant difference between any pair of data (pair of values). In conclusion, there was no reason to continue the experiment using various frequencies and so we chose 130 Hz for further evaluation of cellular movement.

In the second step of the experiment, we used pulsed electric fields with intensities equal to 4, 10 and 20 mA and collected data for control group (0 mV/mm). For the data statistical analysis, we used Kruskal-Wallis test at 0.01 significance level. The results were as follows:

- there were no statistical differences between data on total distance (Fig. 2C), in other words, test confirmed the null hypothesis that all four data samples (control group and three values of pulsed electric field intensity) come from the same distribution; in additional there were no statistical differences between data on straight-line distance between control (0 mV/mm) and experimental data for field values equal to 4 and 10 mV/mm
- there were statistical differences between data on straight-line distance (Fig. 2A); between data on straight-line distance obtained for filed values equal to 4 and 10 mV/mm and data obtained for filed value equal to 20 mV/mm; in addition
- there were statistical differences between data for straightness factor obtained for field values equal to 4 and 10 mV/mm and data obtained for field value equal to 20 mV/mm (Fig. 2B)

**Fig.2.**
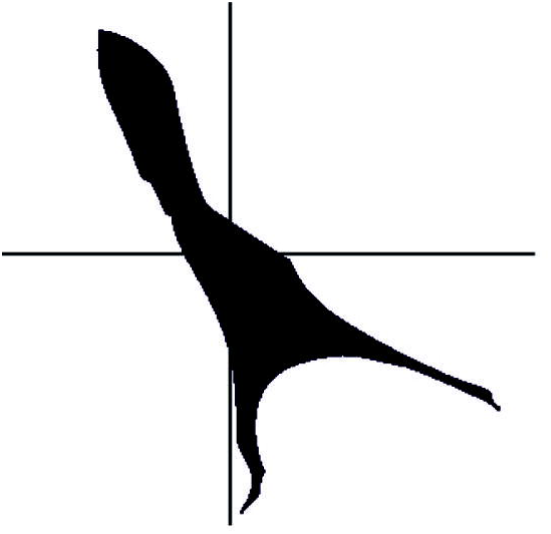
Migration of MSCs: A - straight line distance; B - straightness factor; C - total distance.

Presented figures depict cell images representative for given conditions.

### Influence of pulsed electric field stimulation on the cell glyco-phenotype

We performed a comparative analysis of surface glycosylation in mesenchymal stem cells population after pulsed electric field stimulation, using a panel of plant lectins shown in Table I, known to bind oligosaccharides present in human cells. Staining with the fluorescein-labeled lectins has revealed the dynamics of terminal and internally linked α1,3 and α1,6 Fuc, α1,3 and α1,6 Man, Sia α2,6, and terminal GlcNAc, both on the cell surface, and in intracellular localizations.

Cells from control group were positive for all analyzed lectins. Staining with *Hippeastrum hybrid* (HHL), *Sambucus nigra* (SNA), wheat germ agglutinin (WGA) and *Aleuria aurantia* (AAL) showed glycans located within endoplasmic reticulum and Golgi apparatus (ER/G). Concanavalin A (Con A) and *Narcissus pseudonarcissuss* (NPL) gave the signal both from Golgi apparatus and cytoplasm (cytoplasmic; IC). We observed signals within the different compartments of ER, both in the Golgi vesicles and around the nucleus. Signal closed to the nucleus varied depending on the direction of the electric field. In certain EF conditions, signal appeared highly concentrated forming a very characteristic cap on one side of the nucleus, and called here “polarized ER” (ERp). Similar observation was made by Pu and Zhao (Pu and Zhao 2005). AAL, Con A, SNA and WGA were highlighting multiple cell-cell contact points. Staining for Con A and WGA revealed specific points of cell attachment to the bottom of the culture dishes (glass, bottom substratum). Only in case of these two lectins, the entire cell membrane was outlined by the lectin signals in control cells (cell membrane), but following the EF stimulation, staining was showing only specific membrane domains (cell membrane domains).

To compare the dynamic of complex glycans containing different types of Man linkages, three mannose binding lectin have been used. High amounts of α-mannopyranosyl groups are present as indicated by strong staining with concanavalin A (Con A). Con A recognizes the internal branched Man core trisaccharide (Man α1,6[Man α1,3] Man) of N-linked glycans. HHL pointed the α-mannose residues containing also α-glucosyl structures. Like NPL there appears to be an extended binding site for polymannose structures, not requiring mannose to be at the non-reducing terminus. HHL binds both (α1,3) and (α1,6) linked mannose structures. NPL, isolated from daffodil bulbs, has a specificity toward α-linked mannose, preferring polymannose structures containing (α1,6) linkages. Binding to mannose polymers can occur via internal mannose residues and is not dependent on structural integrity of a non-reducing end sugar. NPL also binds some galactomannans, and differs in other binding characteristics from a related *Galanthus nivalis* lectin.

The presence of poly-*N*-acetyllactosaminyl chains (LacNAc) was clearly demonstrated by the intense staining by wheat germ agglutinin lectin WGA, and the presence of sialic acid was shown by the SNA staining, specific for Neu5Acα2,6Gal/GalNAc.

The orange peel fungal lectin (*Aleuria aurantia* lectin, AAL) recognizes α1,6-linked fucosyl groups occuring on the *N,N'*-diacetylchitobiosyl residues in N-linked glycoproteins (Fukumori et al., 1990).

EF stimulation resulted in the re-distribution of glycans shown by all six lectins (Fig. 3), in both individual signal intensities and localizations. Table II presents key-observations made during analyses of lectin staining patterns of glycan expression and localization.

**Fig.3.**
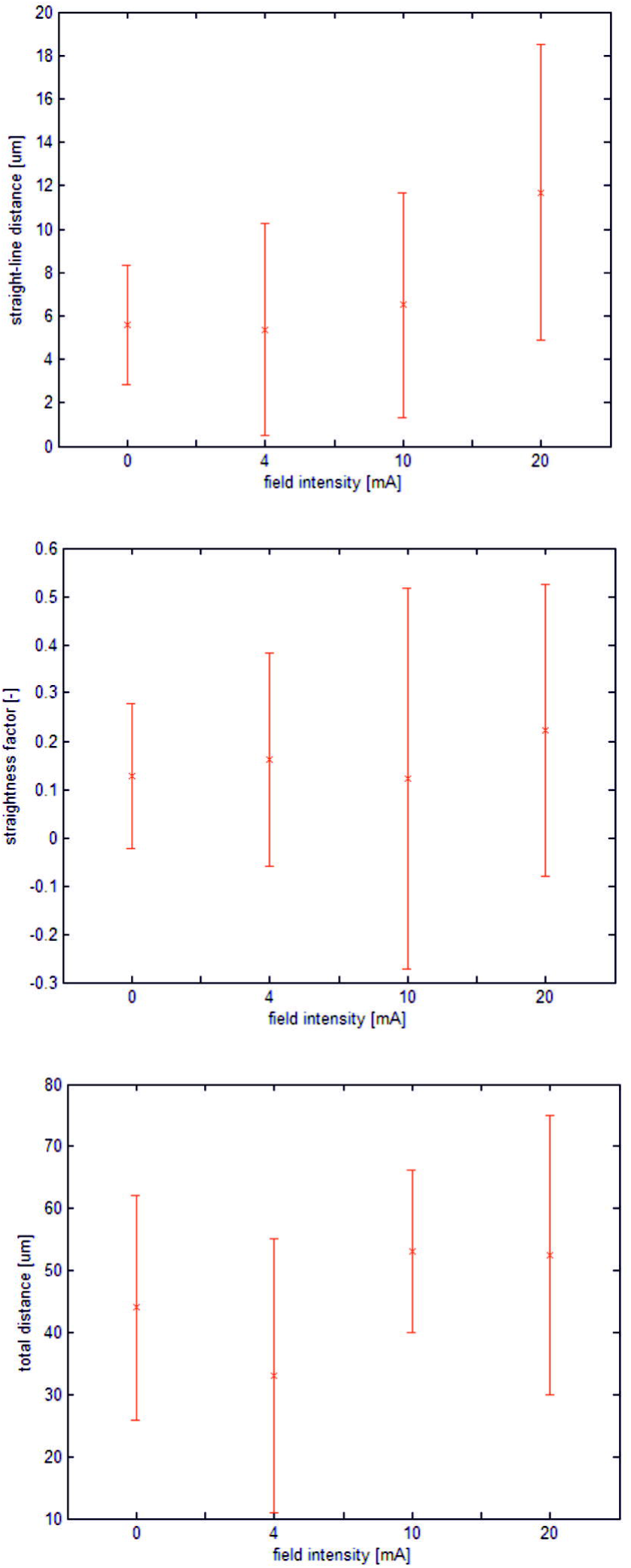
Effect of mannose staining for MSCs after pulsed electric field stimulation by 2 μg/mL fluorescein-labeled FITC-Streptavidin (green) and Cy3-Streptavidyn (red), Hoechst nuclear stain (blue). Scale bar: 10 μm.

**Table 2.**
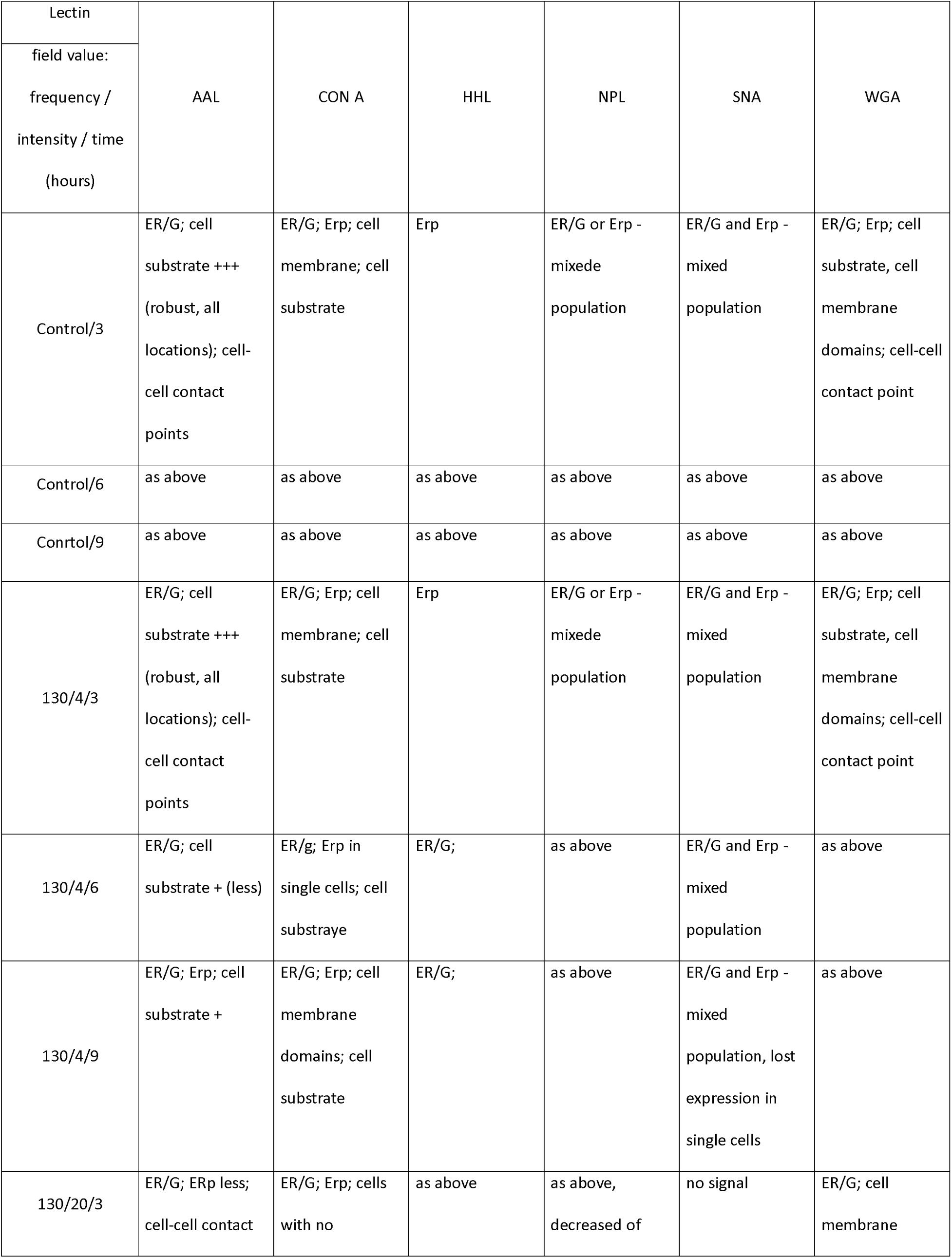
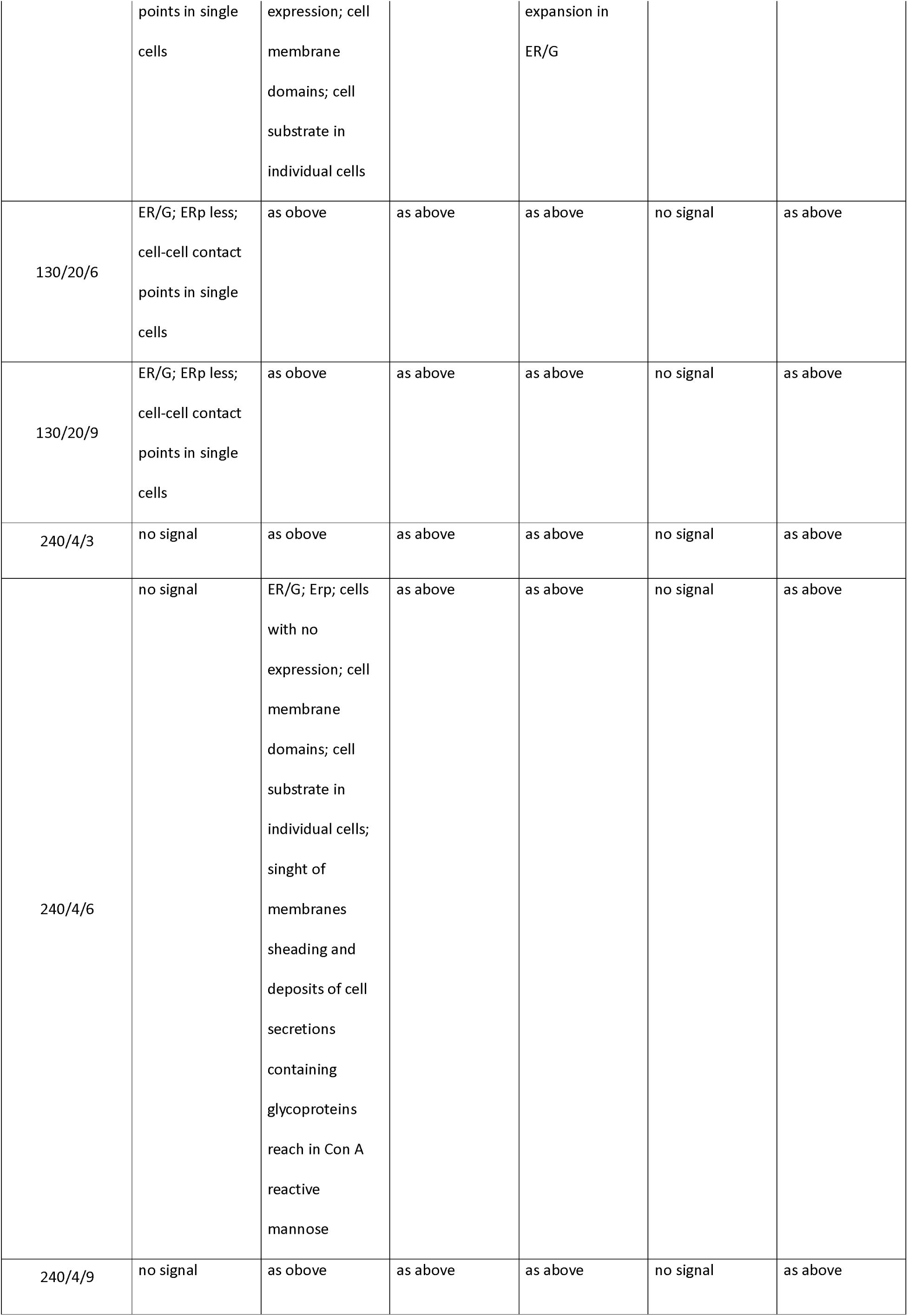
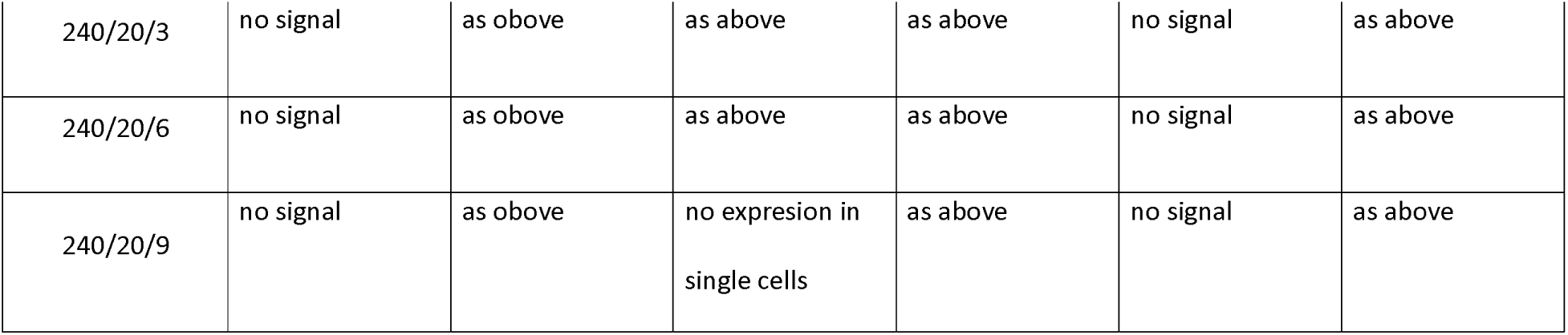
Description of the surface glycans profile changes caused by stimulation of pulsed electric field.

Signals produced by AAL and SNA bindings behaved very similarly. Under exposure to high frequency (240 Hz) EF, target-glycans binding by AAL disappeared, there was no effect of current change on the AAL staining pattern, whereas an increase in the electric field intensity resulted in the loss of SNA signal.

In the case of CON A, very characteristic staining pattern corresponding to membrane shedding, as well as deposits of cell debris and cellular secretions of CON A-reactive materials were observed. In few colonies, there were no visible cells attachment sites away from the main cell population, but the cell-cell contact points were highlighted. This result could be explained by the different types of mannose linkages present in glycans in these locations.

Remodeling of cells surface glycosylation is also visualized by the HHL lectin. Initially visible clear ERp signal disappeared during the long exposure to the low field value and frequency, completely changing cellular localization by moving to the Golgi apparatus. The high frequency electric field resulted in the loss of the HHL signal. We can assume that the polarization of HHL-highlighted ER (ERp) demonstrated polarization of the cell membrane components following cell motion direction.

### Cell viability

To verify that MSCs were not damaged by applied PEFs, the viability of cells following a 3 h, 6 h and 9 h culture exposed to the PEF was compared with that of cells cultured for the same time under similar conditions but without PEF. The viability of cells following a 3 h, 6 h and 9 h culture of stimulation did not differ between groups (*p* > 0.05, one-way ANOVA). This suggests that the application of either low electric field or high electric field within a certain period of time did not result in any measurable damage to the cells.

## Discussion

Aiming at the better understanding of human bone marrow derived mesenchymal stem cells migration and at the development of new techniques to follow the stem cells motility, we analyzed the migration of MSCs in the absence of directional stimuli and in the presence of a pulsed electric field ranging from a low physiological level to the higher than physiological levels, using spinal cord stimulation system. For the first time we have also identified changes in the cell surface glycosylation resulting from stimulation with the pulsed electric field.

Most of the data on the *in vitro* electrotaxis have been obtained by exposing cells to a direct electrical current. In these experiments, metal electrodes directly inserted into the medium, agar or in-house made salt bridges were used (Sato et al., Li and Lin 2012; Cohen et al., 2014; Huang et al., 2013; Song et al., 2007; Wu et al., 2013; Zhao et al., 2011). Effects of alternating electrical current stimulation combined with the direct EF current of very low frequencies have been reported for migration of keratinocytes (Hart et al., 2013).

Despite the successful use of these experimental tools, it is challenging to effectively address more advanced scientific questions and meet higher technical requirements for the rapidly growing electrotaxis research, as well as to the applicability of these protocols in clinical trials. To our knowledge, this is the first work showing the effect of pulsed electric field on the directed migration of MSCs after the application of a dedicated system already used in clinical practice.

We performed stimulation of the MSCs using low and high current and low and high frequencies. In our previous studies (Jezierska-Wozniak et al., 2016) we have shown, that cellular motility increases with the increase of the electric field. In this study we have also analyzed the effect of frequency of electric field on cell motility. We have used two frequencies, 130 Hz and 240 Hz with the highest value of field intensity, i.e. 80 mV/mm, with a 3 hours of observation time to assess the degree of migration. We evaluated statistical differences in the total distance (i.e. distance measured section by section), as well as in the straight-line distance. The Kruskal-Wallis test showed that there was no significant difference between any pair of data. This is in agreement with the results obtained by Jahanshahi and coworkers for electrical stimulation of the progenitor cells in the motor cortex. Furthermore, frequency range does not seem to play any major role in the effectiveness of electrical stimulation on proliferation (Jahanshahi et al., 2013).

We then analyzed three parameters of cell migration such as total distance, distance in straight line and factor of straightness, as well as changes in the profile of cell surface glycans using six plant lectins of the specific glycan-binding properties. The most desirable results, i.e. those in which cells cover the longest possible track in one defined direction, were obtained for the field strength of 20 mA. Under these conditions cells migrated for the long distance and had the highest parameter of straightness.

There are many advantages in using low current for EF stimulation in cells migration studies, where most importantly, excessive heat and unwanted effects on different cell types in the 3D tissue settings are prevented (McCaig et al., 2005; Zhao et al., 2011). Zhao and coworkers (Zhao et al., 2011) have shown that the directional migration occurred at a low threshold and with a physiological EF of ∼25 mV/mm, while increasing the EF-enhanced MSC migratory response. The authors concluded that these conditions selectively directed migration of MSCs while potentially avoiding effects on endothelial cells and their progenitors.

The understanding of biochemical and biophysical changes occurring in cells during migration is rapidly growing. Biochemical studies include genetic manipulation of chemokines and their receptors such as CXCL12/CXCR-4, as well as stimulation of matrix metalloprotease secretion, enhancing chemotaxis and migration of stem cells (Fong et al., 2011; Miller et al., 2008). Studies show also profound changes in the membrane electrical state (Huang et al., 2009) due to the electrostatic membrane surface charge and altered electrodynamic ion fluxes through the membrane channels, resulting in cell membrane permeability changes (Puc et al., 2004) followed by alterations of cytoskeleton structure (Kojima et al., 1992). There is also growing number of reports documenting profile changes of membrane receptors, inter alia epidermal growth factor receptor (EGFR) (Fang et al., 1999) Ca_2_ ions (Huang et al., 2009), and changes in expressions of cytoplasmic signaling proteins such as extracellular signal-regulated kinase (ERK), p38 mitogen-activated kinase (MAPK), cellular sarcoma gene family kinases (Src), protein kinase B (PKB) serine/threonine-specific protein kinase (Akt), and phosphatidylinositol-3 kinase (PI3K) (Zhao et al., 2006).

The ability to recognize distinct carbohydrate determinants turned certain plant lectins into very valuable tools in blood typing (Wu et al., 2009), in evaluating cell differentiation processes (Yura et al., 2008), as cell separation and characterization agents (Naeem et al., 2007), as markers used in microarrays (Tao et al., 2008), in immunological studies, and in studies of cell-cell recognition and cell signaling processes (Kullolli et al., 2008). The importance of glycans in cell therapy has been known since 1988, when Hardy and Tavassoli showed that the carbohydrate moieties present on the stromal cells were important in homing of stem cells bearing the corresponding lectin-like molecules and membrane lectins on a stem cell surface with specificity for galactose and/or mannose-bearing glycoconjugates involved in successful engraftment of hemopoietic progenitor cells in their specific stromal microenvironment (Hardy et al., 1988). Similarly, Hinge and coworkers, showed that the mannose-containing carbohydrate moieties present on the hematopoietic stem and progenitor cells were recognized by mannose-binding lectins and played an important role in the HSPC-protecting effect of these lectins (Hinge et al., 2010). The protective effect of mannose-binding lectins has also been demonstrated by Li and coworkers where the effect of a mannose-binding lectin, NTL, purified from *Narcissus tazetta var. chinensis*, prolonged maintenance and expansion of cord blood CD34+ cells. The results of this study indicate that the effect of NTL on the long term preservation and expansion of early stem /multilineage progenitor cells might be useful in cell therapy strategies for DC34^+^ cells intended for transplantation (Li et al., 2008).

In summary, our result show that: (i) EF value affects the rate of cell migration, in contrast to the EF frequency; (ii) the rate of migration increases with the EF intensity above physiological levels, whereas EF at physiological levels did not affect cell motility, what may be important for the maintenance of the transplanted cells at the lesion site; (iii) EF above physiological levels resulted in changes of the cell surface glycans, whereas physiological EF levels did not change the cell glyco-phenotype as indicated by the lectins used in this study. Since the complex glycosylation of glycolipids and glycoproteins on cell surfaces forms a sophisticated architectural 3D-structure creating the largest cell-surface and cell-substrate adhesion area, changes in this 3D-structure will have a large impact on all properties of cellular adhesion and motility (Amano et al., 2010; Lanctot et al., 2007). Significance and novelty of our approach is based on (i): the recognition of an importance of the dynamics and regulatory function of structural cell surface glycosylation, and (ii) creating (formulating) a novel approach to studies of stem cell membrane glycosylation-based dynamics using a set of lectins binding cell surface carbohydrates.

The success of the stem cell transplantation depends on the effective and functional integration of transplanted cells into the patient's body and into the specific environment of extracellular matrix (ECM) produced and secreted by surrounding native cells. Based on the date results with the targeted cell migration, we used the low value EF remaining within patient’s tolerance range – what is a crucial factor in the cell therapy. Our results show that the application of well-tolerated stimulation parameters should allow securing the transplanted cells at the lesion site, without affecting the phenotype-surface glycans, thus providing a graft that is likely more compatible with the host environment.

If the goal of the stem cell-based therapy is to deliver cells to the lesion site and keeping them there, it seems reasonable to reevaluate the concept of transplanted cells’ homing. Perhaps it is better to choose stimulation / neuromodulation parameters that do not trigger cells to move, but instead keep cells at the lesion site and prevent their migration to other tissues. Therefore, use of the dedicated system for spinal cord stimulation / neuromodulation may provide us with a readily available tool for improving the effectiveness of stem cell therapy.

Use of plant lectins marking specific types of complex glycans allowed us to follow the reprograming of the cell surface glycosylation in response to certain PEF. Here, the specific changes in the cell surface glycans were observed only in cells showing directional movements under treatment with PEF above the physiological level, but in other conditions, such changes may indicate the unwanted migratory potential of cells intended for restorative transplantation.

## Conclusions

The analysis of the stem cells’ glycome dynamics at different stages of differentiation and migration makes possible the exploration of the cell surface glycans as markers of the stem cell functional status, and, in a future – of a transplanted cell – host environment compatibility. Most research focuses on changes in glycan profiles during differentiation of germ cells or induced pluripotent cells (iPS). There is a growing need for better understanding of stem cell functional roles in the selfrenewal and differentiation *in vivo*, as well as through niche interactions and signaling modulation. Since the complex cell surface glycans play a critical role in cell differentiation and migration, we should be aware that the information about glycan dynamics and its regulation may be extensively utilized and likely contribute to the optimizations of the stem cell preparations for future therapeutic applications. For instance, a collective information about the type of transplanted stem cells and the more detailed characterization of their cell surface glycomes using more extensive lectin panel recognizing different 3D glycosylation patterns, can be correlated with the success rate of the transplant and used to optimize protocols for preparations of the therapeutic biomaterial of high quality and efficacy. We believe that our results obtained here using a panel of only six lectins provide a proof of principle, and are an important step in the characterization of stem cells for restorative grafting.

## Competing interests

The authors declare no competing or financial interests.

## Author contributions

Jezierska-Wozniak and Maksymowicz designed and planned the study. Jezierska-Wozniak conducted the study, including cells preparations, cellular motility recordings, data collection and analyses, and analyses of glycosylation dynamics. Barczewska and Grabarczyk performed cell treatment using system for spinal cord stimulation. Lipinski performed evaluation and measurements of the cellular motility. Habich provided support in the statistical data analyses. Jezierska-Wozniak prepared the manuscript draft with the important intellectual input from Maksymowicz, Barczewska and Wojtkiewicz.

## Funding

This work was sponsored by National Science Center (NCN) grants number 2012/07/B/NZ4/01427.

